# Mechanism of DNA-protein crosslink bypass by CMG helicase

**DOI:** 10.1101/2025.03.06.641852

**Authors:** May Thu Kyaw, Sherry Xie, Elias de Lamo Peitz, Hasan Yardimci

## Abstract

DNA-protein crosslinks (DPCs) pose significant barriers to DNA replication and genome integrity. Previous studies have established that bypass of DPCs by the eukaryotic replicative helicase, CMG complex, is critical for DPC repair. We investigated the mechanism of leading-strand DPC bypass by CMG using purified components. Our results reveal that CMG can bypass DPCs in the presence of downstream single-stranded DNA, without the aid of other replisome factors. DPC bypass is a slow process, and its efficiency depends on the size and structure of the protein barrier. DPC bypass does not require CMG to interact with the excluded strand or to unfold the protein adduct, implying that the CMG helicase ring opens to navigate past a DPC. Furthermore, opening of no single MCM interface is essential for bypass, suggesting a flexible ring opening mechanism. Our work highlights the remarkable versatility of the CMG complex in navigating replication challenges, ensuring proper replication fork progression and preserving genome stability.

## INTRODUCTION

In eukaryotic cells, initiation of DNA replication is tightly regulated, and starts with the licensing of replication origins during the G1 phase of the cell cycle. This process involves the loading of the heterohexameric minichromosome maintenance (MCM) 2-7 complexes through the collective actions of origin recognition complex (ORC), CDT1, and CDC6 [1-9]. Double hexamers of the MCM2-7 (hereafter referred to as MCM) complex are assembled around double-stranded DNA (dsDNA) to form pre-replication complexes (preRCs), which remain inactive until cells enter S phase [5, 10-14]. In S phase, Cyclin-Dependent Kinase (CDK) and Dbf4-Dependent Kinase (DDK) are involved in the recruitment of CDC45 and the GINS complex [15-18]. This two-step activation process leads to the formation of the CMG (CDC45/MCM2-7/GINS) complex, the active eukaryotic replicative helicase, which unwinds DNA at the replication fork [19-21]. CMG unwinds DNA using a steric exclusion mechanism, translocating on the leading-strand template in the 3’-to-5’ direction while excluding the lagging-strand template from its central channel [22-26]. Therefore, CMG transitions from encircling dsDNA to single-stranded DNA (ssDNA) during helicase activation, which requires opening an MCM interface to facilitate the process [15, 26, 27]. In addition to unwinding DNA, CMG acts as the platform for replisome components to assemble upon, allowing for leading- and lagging-strand synthesis by polymerases epsilon and delta, respectively [28-34].

During DNA replication, CMG encounters various protein obstacles such as histones, cohesins, and RNA polymerases [35, 36]. While such nucleoprotein complexes may be disrupted, CMG can also encounter covalent DNA-protein crosslinks (DPCs), which often require an involved proteolysis mechanism for their removal [37-39]. DPCs can be induced by exogenous factors such as exposure to certain chemicals and chemotherapeutic agents, radiation, as well as endogenous enzymatic processes [40-43]. Whilst lagging-strand DPCs do not significantly affect CMG progression [23], DPCs found on the leading-strand template may block the replicative helicase and subsequent synthesis machinery – potentially leading to fork collapse. In the seminal work from the Walter laboratory, it was demonstrated that CMG could bypass a leading-strand DPC in *Xenopus* egg extracts with the help of an accessory helicase, RTEL1 [38]. RTEL1, a 5’ to 3’ DNA helicase, has been shown to unwind DNA downstream of a DPC to create ssDNA adjacent to the adduct and thereby promote DPC traversal by CMG [38, 44]. This bypass mechanism facilitates the recruitment of SPRTN metalloprotease to enable subsequent DPC proteolysis [37, 38].

The mechanistic details of how the CMG helicase bypasses leading-strand DPCs, while encircling this strand, are not fully understood. It is speculated that CMG either denatures protein adducts through its central channel or opens its MCM ring to accommodate the intact protein adduct during bypass. We previously showed another replicative helicase, SV40 large T-antigen (LTag) bypasses DPCs on the translocation strand likely by opening its homohexameric ring structure [45]. While it is established that MCM interfaces open during helicase loading and activation, it is unclear whether CMG can similarly open its ring while actively unwinding DNA. Furthermore, it remains uncertain if DPC traversal by CMG, as observed in egg extracts, requires accessory factors beyond RTEL1. Elucidating the details of this mechanism is essential to understand the intricate interplay between eukaryotic DNA replication and repair processes.

To unravel the mechanism by which the CMG helicase bypasses bulky protein adducts on the leading-strand template, we conducted DNA unwinding assays using recombinant CMG with a variety of model fork DNA substrates. Our experiments showed that purified CMG can navigate past a leading-strand adduct when ssDNA is present downstream, eliminating the need for additional replisome factors, apart from RTEL1. Additionally, our data indicates that CMG’s interaction with the excluded DNA strand is not critical for DPC traversal. Our findings also reveal that the bypass of DPCs by CMG does not depend on the denaturation of the protein adduct, implying that CMG facilitates bypass by opening its MCM ring. Furthermore, by experimentally inducing dimerization of adjacent MCM subunits, we determined that the opening of any individual interface is not a prerequisite for bypass indicating the ability of opening multiple interfaces within the MCM ring.

## RESULTS

### CMG can bypass a leading-strand DPC when presented with downstream ssDNA

To better understand the mechanism by which CMG traverses a leading-strand DPC, we sought to identify the minimal components required for this process. To this end, we used a single-turnover DNA unwinding assay, previously described in Kose et al. [46], containing recombinantly purified *Drosophila melanogaster* (*Dm*) CMG and synthetic fork DNA substrates containing a HpaII methyltransferase (M.HpaII) covalently crosslinked at specific sites [47]. For efficient CMG loading, all fork substrates were designed to have a 40-nucleotide (nt) 3’ polyT tail. Moreover, the 5’ tail was designed with GAGC repeats that form secondary structures, serving a dual function: preventing CMG from binding to the 5’ tail and ensuring that CMG, once bound to the 3’ tail, remains active at the fork junction [46]. To perform unwinding assays, CMG was initially bound to the fork DNA with ATPγS, an ATP analogue that precludes DNA unwinding by *Dm*CMG [46], followed by ATP addition to trigger unwinding (Figure 1A). To hinder the reannealing of separated strands, oligonucleotides complementary to the excluded strand were added alongside ATP. Furthermore, excess polyT oligonucleotides were included to prevent CMG reengagement with fork DNA [46]. The presence of a Cy5 fluorophore on the excluded strand enabled differentiation between unwound and intact DNA via native polyacrylamide gel electrophoresis (PAGE). In line with our previous observations [22], we found that CMG was able to efficiently unwind non-adducted fork DNA, while a leading-strand M.HpaII adduct (MH^Lead^) hindered CMG-mediated unwinding (Figure 1B) indicating that CMG is unable to bypass MH^Lead^.

**Figure 1.**
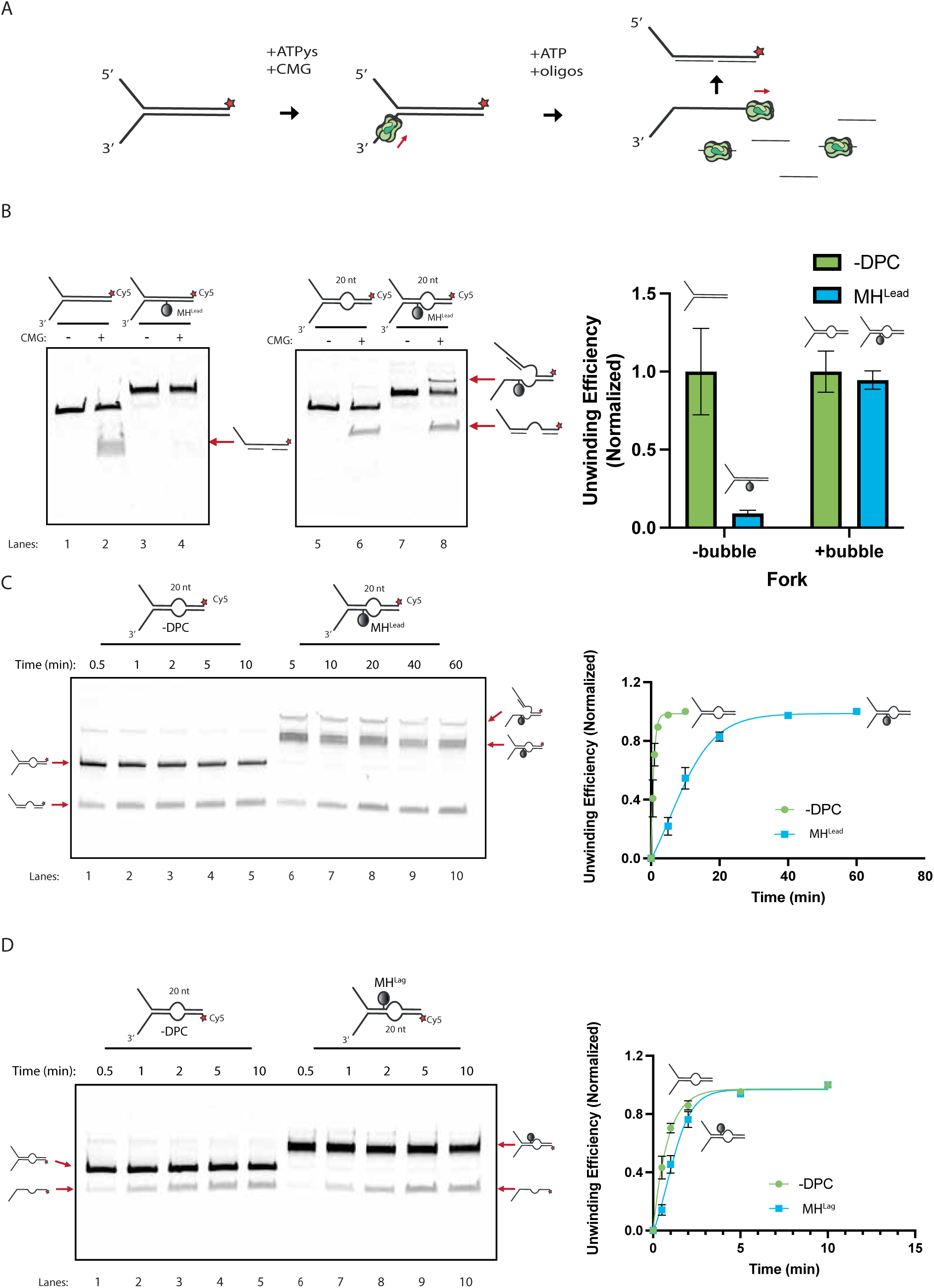
CMG can bypass a leading-strand DPC when presented with downstream ssDNA. **(A)** Diagram showing the method used for measuring single-turnover DNA unwinding by pre-loaded CMG onto a model fork DNA substrate. **(B)** Unwinding of fork DNA with and without a MH^Lead^ in the presence or absence of a downstream 20-nt ssDNA bubble. Fork DNA substrates contained a 3’ Cy5 fluorophore on the non-translocation strand, which did not contain DPC. CMG was pre-bound to fork substrates in the presence of ATPγs at 37°C for 1 hour. ATP and trap oligos were later added to initiate unwinding. Reactions were further incubated at 30°C for 1 hour before stopping with SDS. DNA was subsequently separated on an 8% native PAGE and Cy5 fluorescence was imaged. The right panel displays the quantification of unwinding efficiency, normalized with respect to the unwinding efficiency on non-adducted fork DNA. The data represent mean ± SD from three independent experiments. **(C-D)** Time-course unwinding assay on fork DNA containing either (C) MH^Lead^ or (D) MH^Lag^. Both forks contained a 5’ Cy5 fluorophore on the non-translocating strand and a 20-nt ssDNA bubble downstream of the DPC. Quantification of normalized unwinding efficiencies are shown on the right panels, with data represented as the mean ± SD from three independent experiments.

Previous work by Sparks et al. [38] demonstrated that RTEL1 is essential for generating ssDNA downstream of a DPC, facilitating the traversal of the adduct by the CMG helicase in egg extracts. Notably, this study also showed that a pre-introduced ssDNA bubble adjacent to the DPC could substitute for RTEL1 activity, enabling CMG to bypass the DPC. To investigate whether the presence of ssDNA downstream alone is sufficient for CMG to bypass a DPC, we engineered a fork DNA substrate to incorporate a 20-nt ssDNA bubble, which included a polyT sequence on both strands, placed 7 base pairs (bps) away from the base on which MH^Lead^ is crosslinked. CMG demonstrated efficient unwinding of this modified fork containing the ssDNA bubble, even when the MH^Lead^ was present (Figures 1B and 1C). This data suggests that the CMG helicase, once loaded onto the fork DNA’s 3’ polyT tail, can unwind past a leading-strand DPC. To demonstrate that CMG bypasses the DPC, instead of a second CMG binding the 20-nt DNA bubble, we designed a fork substrate without the 3’ fork branch, but containing the same 20-nt ssDNA bubble (Figure S1). Additionally, we introduced a Cy5-labeled oligonucleotide downstream of the ssDNA bubble to detect CMG activity starting from the bubble. The absence of the 3’ tail resulted in no strand displacement, demonstrating that CMG does not bind to the 20-nt bubble (Figure S1). This confirms that a downstream 20-nt ssDNA bubble is sufficient for the CMG helicase to bypass a MH^Lead^ adduct.

In *Xenopus* egg extracts, replisomes paused for 15 minutes on average when encountering an MH^Lead^ before complete traversal [38]. The TIM-TIPIN complex (Csm3-Tof1 complex in yeast) is known to induce temporary pausing at various replication fork barriers in cells [48], which could explain replisome stalling in extracts after collision with a leading-strand DPC. To assess if the pause at a DPC is a characteristic feature of the CMG helicase itself or due to other replisome factors, we performed a time-course unwinding assay on the fork substrate with a 20-nt ssDNA bubble, both with and without the MH^Lead^. The unwinding of the non-adducted fork reached a plateau at 5 minutes, whereas the fork with the MH^Lead^ took over 20 minutes to reach a similar extent of unwinding, mirroring the delay seen in egg extracts (Figure 1C). Moreover, this delay was specific to leading-strand DPCs as evidenced by the fact that a lagging-strand M.HpaII adduct (MH^Lag^) only caused a brief delay (Figure 1D). Thus, pausing at a leading-strand DPC is an intrinsic behaviour of the CMG helicase, independent of other replisome components.

Given that the full unwinding of the adducted fork substrate required an extended duration, we considered the possibility that CMG might unwind DNA to the site of the DPC, with the downstream DNA melting gradually over time rather than being actively unwound by CMG. To determine whether CMG’s helicase activity is crucial for unwinding the duplex DNA region past the DPC, we stopped the CMG helicase activity around the moment they encounter the DPC (10 minutes), by introducing a 10-fold excess of ATPγS over ATP (Figure S2A). If CMG’s ATPase activity is essential for unwinding past the DPC, the introduction of excess ATPγS should preserve the intermediate product, where only the first half of the fork DNA has been unwound. Conversely, if the downstream DNA were simply melting, we would expect to see an increase in completely unwound DNA and a decrease in the intermediate species over time, regardless of ATPγS addition. Adding excess ATPγS stabilized the intermediate species and prevented further unwinding of the DNA (Figures S2B and S2C), confirming that CMG’s ATPase activity is necessary to unwind DNA beyond a DPC. Our findings collectively demonstrate that CMG can bypass leading-strand DPCs when ssDNA is present downstream. This is in line with the known role of the accessory helicase RTEL1 in facilitating DNA unwinding past a DPC in egg extracts. Furthermore, these results support the notion that CMG does not rely on other replisome components other than an accessory helicase to bypass bulky adducts.

### CMG does not denature protein roadblocks during bypass

We considered the possibility that the CMG helicase bypasses leading-strand DPCs by denaturing protein roadblocks within its central channel. While some helicases, such as FANCJ, have been shown to denature proteins crosslinked to DNA [49], isolated CMG does not exert as much force while unwinding DNA as other replicative helicases, for instance L-Tag, which can disrupt even strong biotin-streptavidin interactions [45]. Therefore, it seemed plausible that CMG might bypass DPCs without denaturing them. To test this hypothesis, we performed a single-turnover unwinding assay with fork DNA where monomeric streptavidin (mSA) was attached to a biotin on the translocation strand adjacent to a ssDNA bubble (Figure 2A). As the interaction between mSA and biotin is structurally dictated, if CMG denatures the mSA during bypass, it will disrupt this interaction and release mSA from the DNA. Excess free biotin was added with ATP to capture any dissociated mSA and prevent its rebinding to the DNA (Figure S3). Following unwinding of the fork by CMG, biotinylated strand remained bound to mSA suggesting that CMG does not denature leading-strand protein roadblocks during bypass (Figure 2B). Therefore, we conclude that CMG does not denature leading-strand DPCs but instead traverses them by opening its MCM ring, allowing the helicase to accommodate the bulky protein adduct in a manner that circumvents steric hindrance.

**Figure 2.**
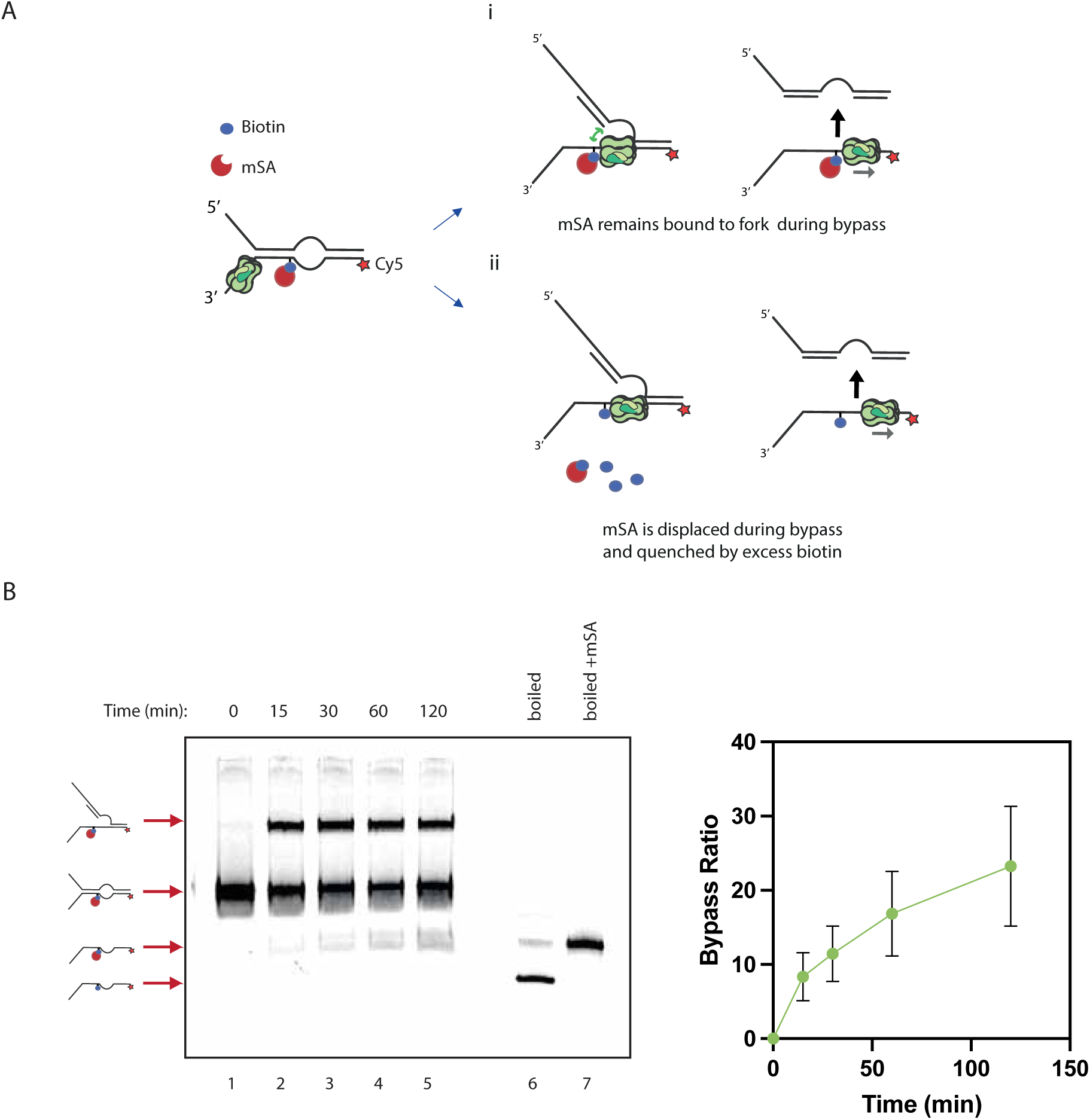
CMG does not denature protein roadblocks during traverse. **(A)** Cartoon model showing potential scenarios regarding how CMG bypasses monomeric streptavidin (mSA) bound to biotin on the translocation strand. To follow mSA displacement, the biotin-modified translocation strand was labeled with Cy5. Monomeric streptavidin either (i) remains bound to the biotin on DNA or (ii) is displaced by CMG. Excess biotin quenches the displaced mSA in (ii). **(B)** After pre-binding mSA to biotin-modified fork DNA, CMG was bound to fork with ATPγs. ATP, trap oligonucleotides, and excess biotin were added to initiate unwinding. Samples were withdrawn at various time points and reaction was stopped with SDS. Samples were then separated on PAGE and Cy5 fluorescence was imaged. The right panel represents the quantification of the bypass ratio of unwound product divided by the sum of the intermediate and unwound products. The data are represented as the mean ± SD from three independent experiments. Although the overall efficiency of biotin-bound mSA bypass was lower compared to MH^Lead^ bypass, our assay detected only mSA-bound unwound DNA, with no evidence of biotin-modified strand devoid of mSA.

### 5-nt ssDNA is sufficient for DPC traversal by CMG

We envisaged two possible models to explain how the CMG complex might bypass leading-strand DPCs by opening its MCM ring. The first scenario involves CMG employing a ‘hopping’ mechanism, wherein upon encountering the DPC, the helicase temporarily disengages from the DNA and then rebinds downstream onto the ssDNA bubble (Figure 3A, left). However, this seems implausible based on our findings that CMG cannot bind *de novo* to a 20-nt ssDNA bubble (Figure S2). Additionally, in our experiments, the presence of excess polyT oligos would likely inhibit any CMG helicase that had detached from the DNA. Thus, we considered a second mechanism where the helicase could partially open its ring and move over the adduct incrementally, in a manner akin to ‘crawling’ (Figure 3A, right).

**Figure 3.**
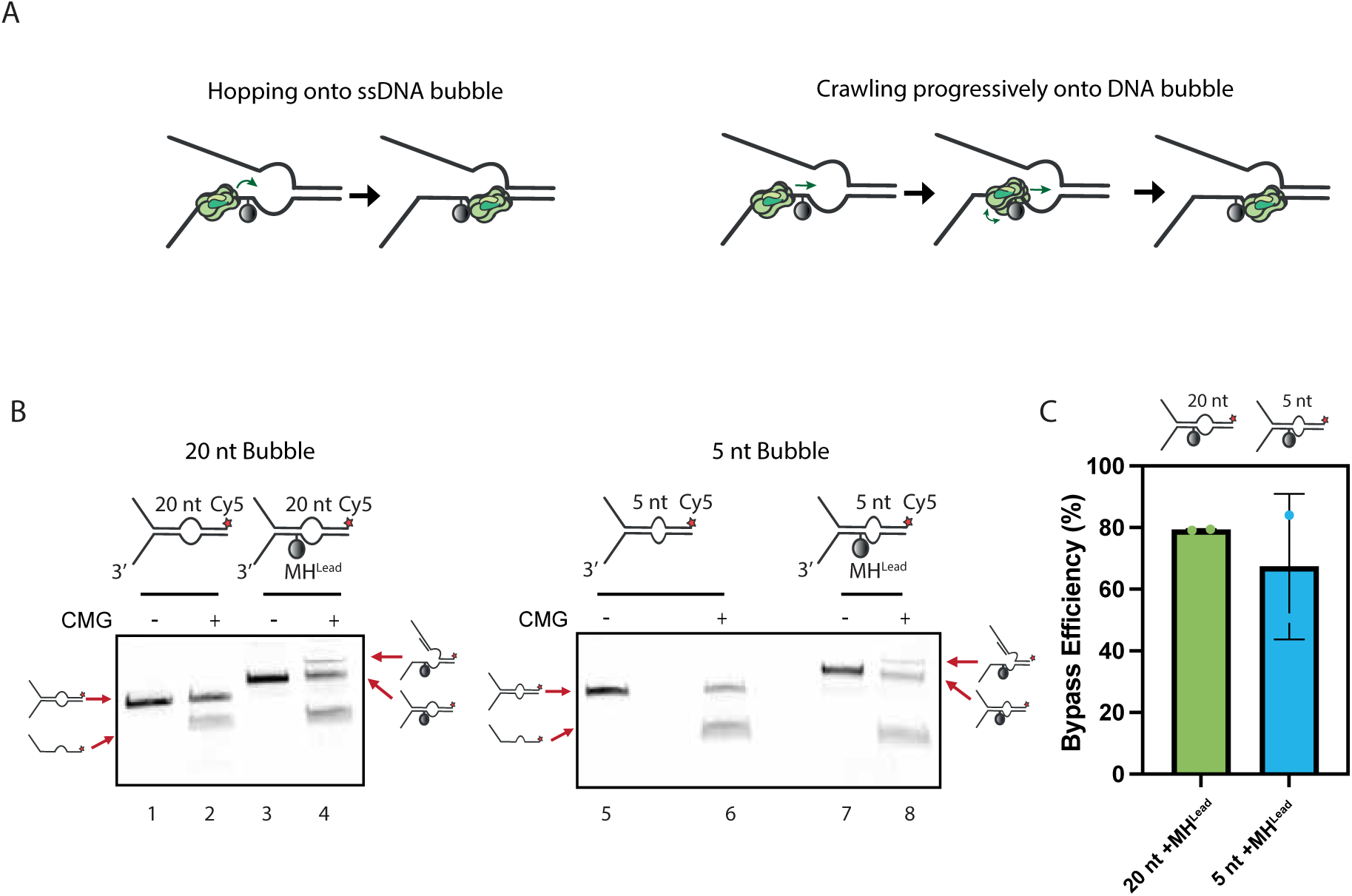
Short downstream ssDNA is sufficient for DPC traversal by CMG. **(A)** Cartoon diagram illustrating possible bypass mechanisms on how CMG uses downstream ssDNA. The left panel shows CMG ‘hopping’ onto the preformed downstream ssDNA in a concerted fashion. This scenario implies that ssDNA must be long enough for CMG to bind. The right panel illustrates a scenario where CMG ‘crawls’ slowly over the adduct, grasping the downstream ssDNA. In this scenario, the downstream ssDNA does not to necessarily need to be as long as CMG’s footprint. **(B)** Unwinding assays on fork DNA substrates with different (left) 20-nt or (right) 5-nt ssDNA with or without MH^Lead^. Fork DNA substrates contained a 3’ Cy5 fluorophore on the non-adducted excluded strand. **(C)** Quantification of the CMG-driven unwinding efficiency on MH^Lead^-crosslinked fork substrates containing either 20- or 5-nt ssDNA bubble, with the data represented as the mean ± SD from two independent experiments.

The combination of 20-nt ssDNA bubble and the 7-nt DNA between MH^Lead^ and the bubble utilized in our prior experiments approximates the 20- to 40-nt footprint of CMG [26]. If CMG were to hop, it would need at least ssDNA similar to its footprint to reattach downstream of the DPC. A crawling mechanism, by contrast, could operate with less ssDNA than the helicase’s footprint. We thus decreased the ssDNA bubble to 5 nt, substantially smaller than CMG’s footprint, to test this hypothesis. Remarkably, CMG still efficiently traversed the MH^Lead^ on this substrate (Figures 3B and 3C), suggesting that it does not leap over the DPC, but instead engages with the available ssDNA and inches over the adduct.

### Interactions between CMG and the excluded DNA strand are dispensable for bypass

Previous structural studies have identified an interaction between the CMG helicase and the excluded DNA strand at the replication fork, which has been suggested to shield the helicase from ubiquitination and disassembly during fork progression [22, 25, 50]. Given the potential for the CMG helicase ring to open while navigating past DPCs, we investigated whether this interaction with the excluded strand is necessary to maintain the helicase on DNA. Specifically, we posited that the excluded strand might act as a guide rail, enabling CMG to move past a leading-strand DPC while also being safeguarded from ubiquitination.

To investigate the necessity of CMG’s engagement with the excluded strand for DPC traversal, we constructed a fork DNA substrate featuring a ssDNA gap with a DPC positioned on the ssDNA. To this end, we crosslinked protein G to fork DNA, adopting a protocol described in a previous study [51]. The non-translocation strand was designed with two sections: the first containing a fluorescein label, and the second, downstream of the ssDNA gap, carrying a Cy5 label (Figure 4A).

**Figure 4.**
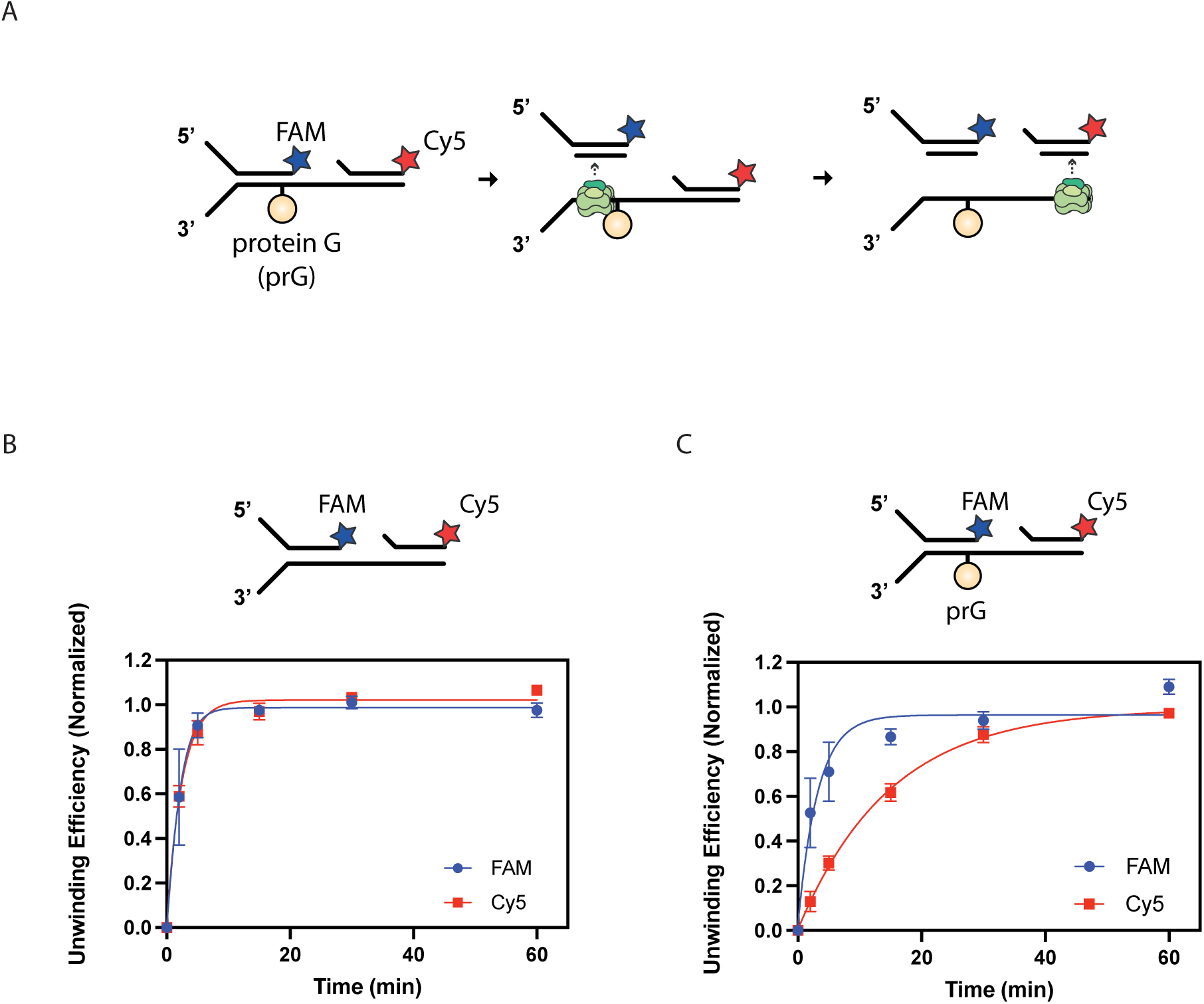
Interactions between CMG and the excluded are dispensable for bypass. **(A)** A cartoon diagram of the experiment. The fork DNA substrate contained a ssDNA gap of 10 nt with or without a covalent leading-strand protein G crosslink (prG^Lead^) placed on the ssDNA gap. The first half of the fork carried a 3’ fluorescein amidite (FAM), and the oligonucleotide downstream of the ssDNA gap contained a 3’ Cy5 fluorophore. Both labeled strands were engineered to form hairpins on their 5’ ends, blocking CMG binding and promoting their displacement by CMG. **(B-C)** Quantification of displacement efficiency of FAM- and Cy5-labeled strands on fork DNA (B) without DPC and (C) with prG^Lead^. The data is represented as the mean ± SD from three independent experiments.

On this fork substrate, when CMG unwinds the DNA up to the leading-strand protein G crosslink (prG^Lead^), it would displace the fluorescein-labeled oligonucleotide. This displacement results in the CMG losing its interaction with the excluded strand. Subsequently, if the CMG can also displace the Cy5-labeled oligonucleotide, it would demonstrate its ability to bypass a leading-strand DPC without the need to engage with the excluded strand. In experiments conducted on a fork substrate without any adduct, CMG displaced both labeled strands within 5 minutes (Figures 4B and S4). When a prG^Lead^ was introduced to the fork, the Cy5 strand was still efficiently displaced (Figures 4C and S4). However, while the fluorescein-labeled oligonucleotide was displaced within 5 minutes, the displacement of the Cy5-labeled oligonucleotide reached a plateau at around 30 minutes (Figure 4C), mirroring the delay observed with substrates containing MH^Lead^ and ssDNA bubble (Figure 1C). These findings demonstrate that CMG can bypass a leading-strand DPC without relying on interactions with the excluded strand, implying that its retention on DNA is facilitated through interactions solely with the translocation strand.

### Size and structure of the protein barrier affect bypass efficiency

Diverse DNA-binding proteins can crosslink to DNA within cells, creating a broad array of DPCs. Our experiments have demonstrated that the CMG helicase can bypass a HpaII methyltransferase (40 kDa) crosslinked to DNA via an internal amino acid, as well as a protein G (26 kDa) attached at its N terminus. To explore whether the specific characteristics of DPCs influence CMG’s bypass ability, we introduced various recombinant proteins crosslinked to fork DNA. The CMG helicase effectively unwound DNA when encountering protein A/G (59.7 kDa) — structurally similar to protein G but significantly larger (Figure 5). Yet, it failed to bypass the tetrameric wild-type streptavidin (SA), which has a molecular weight comparable to protein A/G but differs markedly in structure (Figure 5). To discern if the size or structure of SA hindered CMG, we crosslinked mSA (15 kDa), a structurally similar but smaller SA variant. CMG successfully unwound DNA with the mSA crosslink. This suggests that CMG may need to expand its MCM ring more extensively to navigate past SA than it does for protein A/G or mSA. The data, therefore, imply that both the size and structure of the protein adduct affect CMG’s capability to bypass a DPC.

**Figure 5.**
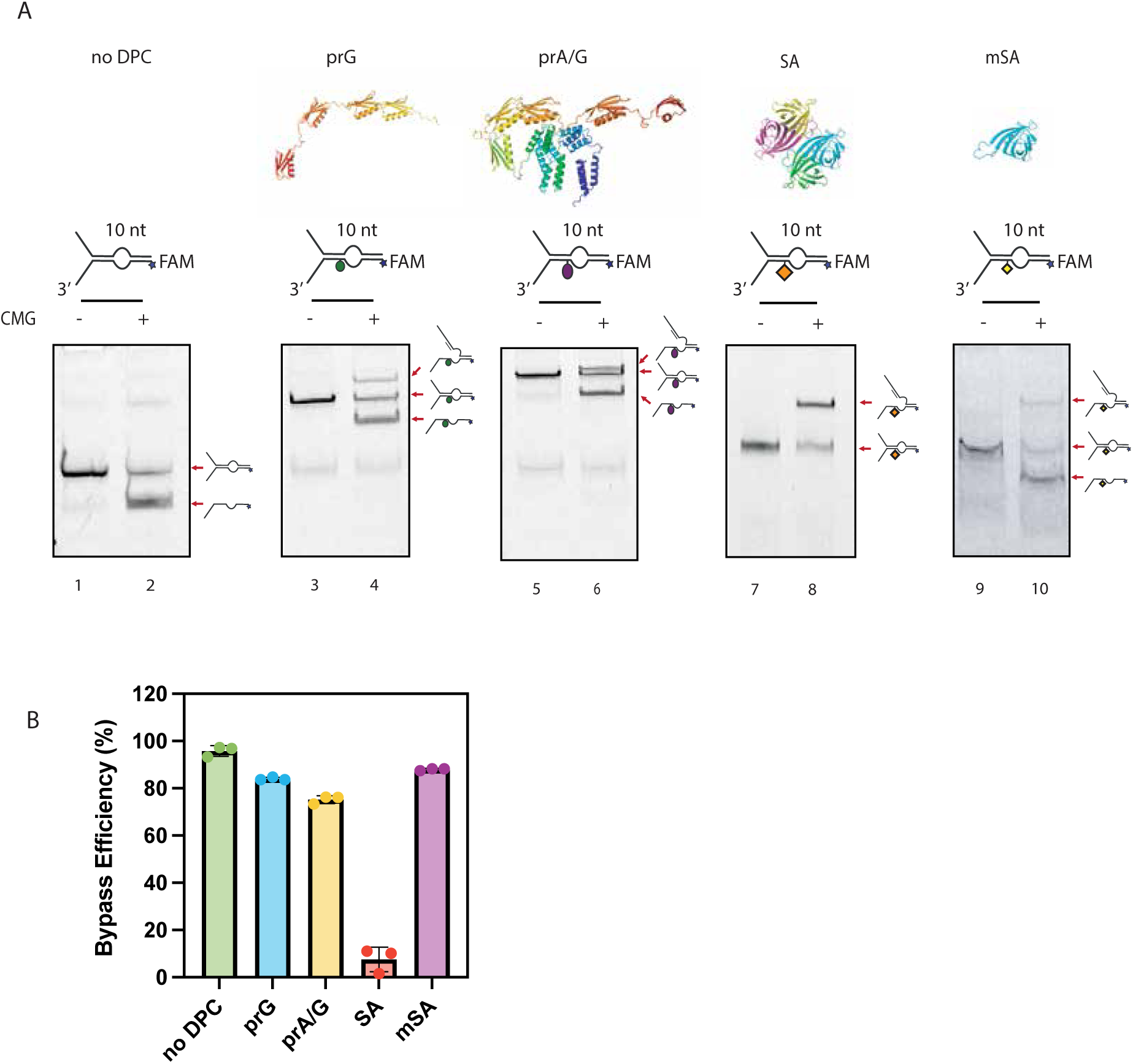
Size and structure of the protein barrier affect bypass efficiency. **(A)** Single turnover CMG-driven unwinding of fork DNA containing various proteins crosslinked to the translocation strand through thiol-modified oligonucleotides. Forks contained 10-nt ssDNA bubble following the DPC and were labeled with fluorescein amidite (FAM) on the translocating strand. AlphaFold predictions of protein G (prG), protein A/G (prA/G) as well as previously solved structures of streptavidin (SA, PDB:6J6J) and monomeric streptavidin (mSA, PDB:4JNJ) are shown. **(B)** Quantification of DPC bypass efficiency on fork substrates containing no DPC, prG, prA/G, SA, or mSA as leading-strand DPCs. The data are represented as the mean ± SD from three independent experiments.

### CMG can open more than one MCM interface to traverse DPCs

Given the precise regulation of CMG helicase assembly, where the MCM 2/5 interface is shown to be critical during MCM loading [8, 52, 53], we explored whether the bypass of leading-strand DPCs by CMG necessitates opening a unique MCM interface. To probe the role of MCM interfaces, we engineered a series of CMG variants, each with two adjacent MCM subunits modified to control the closure of their interface using an inducible system (Figure S5A). This was achieved using the FK506 binding protein (FKBP) and the FKBP-rapamycin binding (FRB) domain of the mammalian target of rapamycin (mTOR) kinase, which dimerize in the presence of rapamycin [54]. Employing this system, which has been previously used to establish the critical nature of the MCM 2/5 interface in the MCM complex’s loading onto DNA [52], we assessed the DPC bypass efficiency of each CMG construct with and without rapamycin. None of the mutant CMG constructs showed a decline in bypass efficiency when rapamycin was introduced, suggesting that the opening of a unique MCM interface is not requisite for bypassing a leading-strand DPC (Figure 6). However, it was not possible to prove dimerization of FRB and FKBP for the entirety of an unwinding assay. To circumvent this issue, we covalently crosslinked adjacent MCM subunits by using DogCatcher and DogTag, a peptide-peptide conjugation strategy [55]. To covalently close the MCM 2/5 interface, we employed an inducible SnoopLigase-mediated system [56, 57]. This method led to 50-80% crosslinking of neighbouring MCM subunits (Figures S5B and S5C). Importantly, none of the covalently closed CMG variants showed decline in DPC bypass efficiency (Figures S5D-S5G). Together, our results imply that CMG does not rely on the opening of a single MCM interface to bypass DPCs and may instead open multiple interfaces.

**Figure 6.**
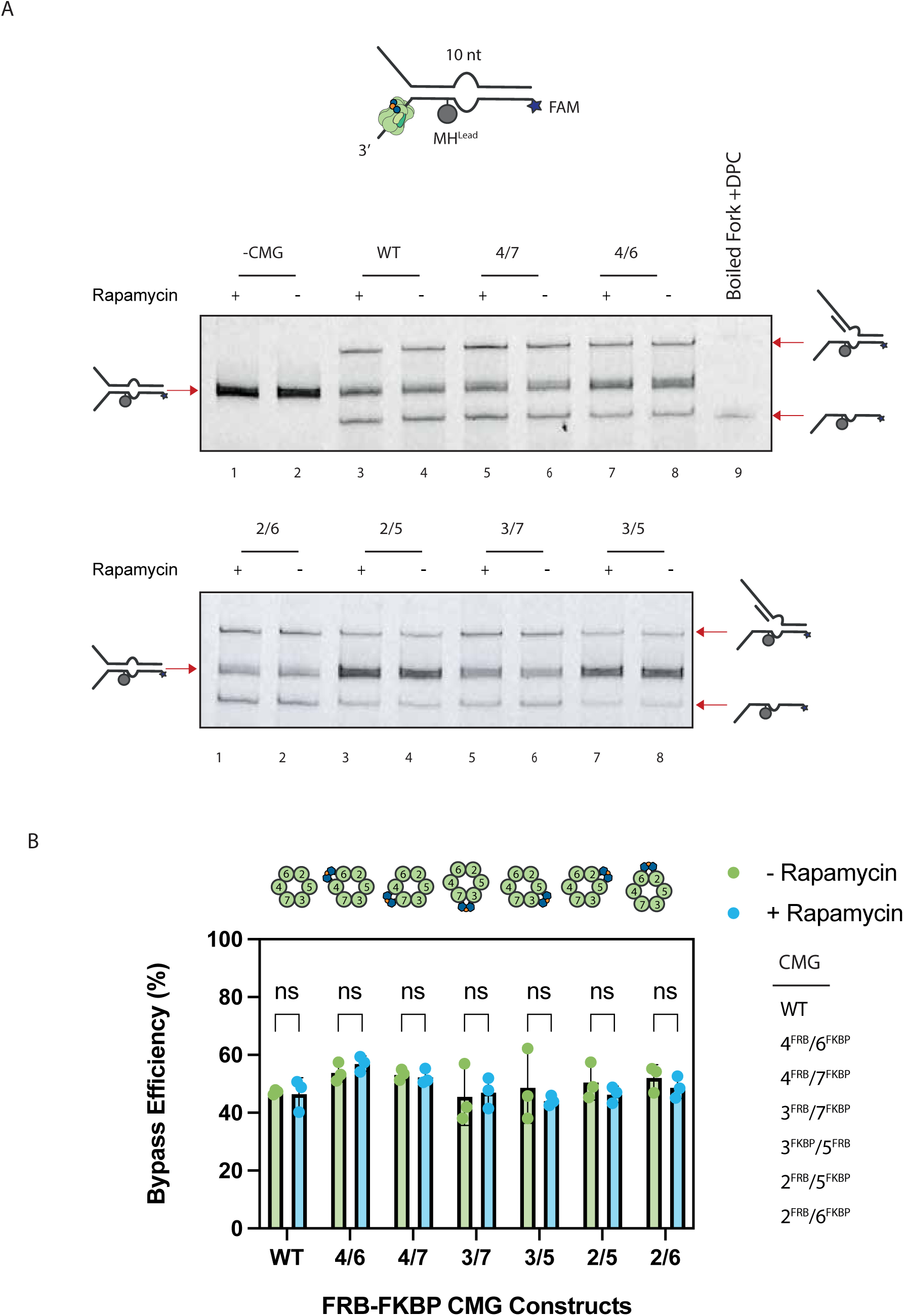
CMG can open more than one MCM interface to traverse DPCs. **(A)** Single-turnover fork DNA unwinding assay with various FRB-FKBP CMG constructs in the absence or presence of rapamycin. The fork substrate contained MH^Lead^ followed by a 10-nt ssDNA bubble and fluorescein amidite (FAM) on the translocating strand. CMG was pre-bound to fork substates in the presence of ATPγs. Samples were halved and incubated for an additional 10 minutes in the absence or presence of 1 µM rapamycin. ATP and trap oligonucleotides were then added to initiate unwinding. Reactions were further incubated at 30°C for 1 hour and stopped with SDS. DNA was then separated on native PAGE and FAM fluorescence was imaged. **(B)** Quantification of DPC bypass efficiency of the different CMGs in the absence and presence of rapamycin. We quantified the ratio of fully unwound products to intermediates where DNA is unwound only up to the DPC. Data represented as the mean ± SD from three independent experiments.

## DISCUSSION

Our study provides new insights into the mechanisms of DPC bypass by the CMG helicase, an essential process for maintaining genome stability during DNA replication. CMG is capable of navigating past leading-strand DPCs provided there is downstream ssDNA, without the necessity to denature the protein obstacle suggesting that MCM ring opening facilitates DPC bypass. Time-course analyses indicate that while DPCs slow CMG’s movement, they do not completely halt its progress. Additionally, our data reveal that CMG’s interaction with the excluded strand is dispensable for DPC traversal. The ability of CMG to bypass DPCs appears to be influenced by the size and structure of the crosslinked protein. Importantly, our experiments show that CMG can traverse DPCs even when each of the interfaces between the six MCM subunits is individually closed, implying the helicase’s bypass ability does not depend on the opening of any specific MCM interface.

### A conserved mechanism for traversal of bulky obstacles by CMG helicase

CMG may employ a conserved mechanism to traverse leading-strand barriers that cannot be accommodated within its central channel. This is exemplified by previous work in *Xenopus* egg extracts, which showed that G4 structures on the leading-strand template led to CMG stalling. Subsequently, the DEAH-box helicase 36 (DHX36) unwinds past the G4 structures to facilitate bypass by CMG [58], a process reminiscent of RTEL1’s role in DPC bypass [38]. This consistency across different types of obstacles hints at a fundamental mechanism CMG uses to bypass large barriers, likely relying on the presence of ssDNA downstream from the barrier.

We propose a mechanistic model wherein CMG initially stalls upon encountering a DPC or other stable barriers on the translocation strand (Figure 7). Following this encounter, an accessory helicase, acting on the opposite strand to CMG, unwinds the DNA, creating ssDNA beyond the barrier that is essential for bypass. This unwinding does not require the accessory helicase to be highly processive; it only needs to generate a small stretch of ssDNA. This action may be a repetitive process, with the accessory helicase unwinding and dissociating, and the fork DNA potentially reannealing up to the barrier until CMG successfully bypasses it. We speculate that an interface in the N-terminal domain (NTD) of the MCM complex, positioned at the forefront, may be the first to open. This could happen stochastically or be induced by the physical pressure of CMG against the barrier. Following the NTD opening, the C-terminal motor domain (CTD) would advance, moving CMG forward. Once the NTD clamps onto the downstream ssDNA, the C-terminal motor domain (CTD) is then primed to open. This allows the helicase to continue its progression past the barrier, either through active translocation along the DNA or diffusion facilitated by the NTD on the ssDNA beyond the roadblock. After the CTD clears the barrier, CMG can resume unwinding the replication fork.

**Figure 7.**
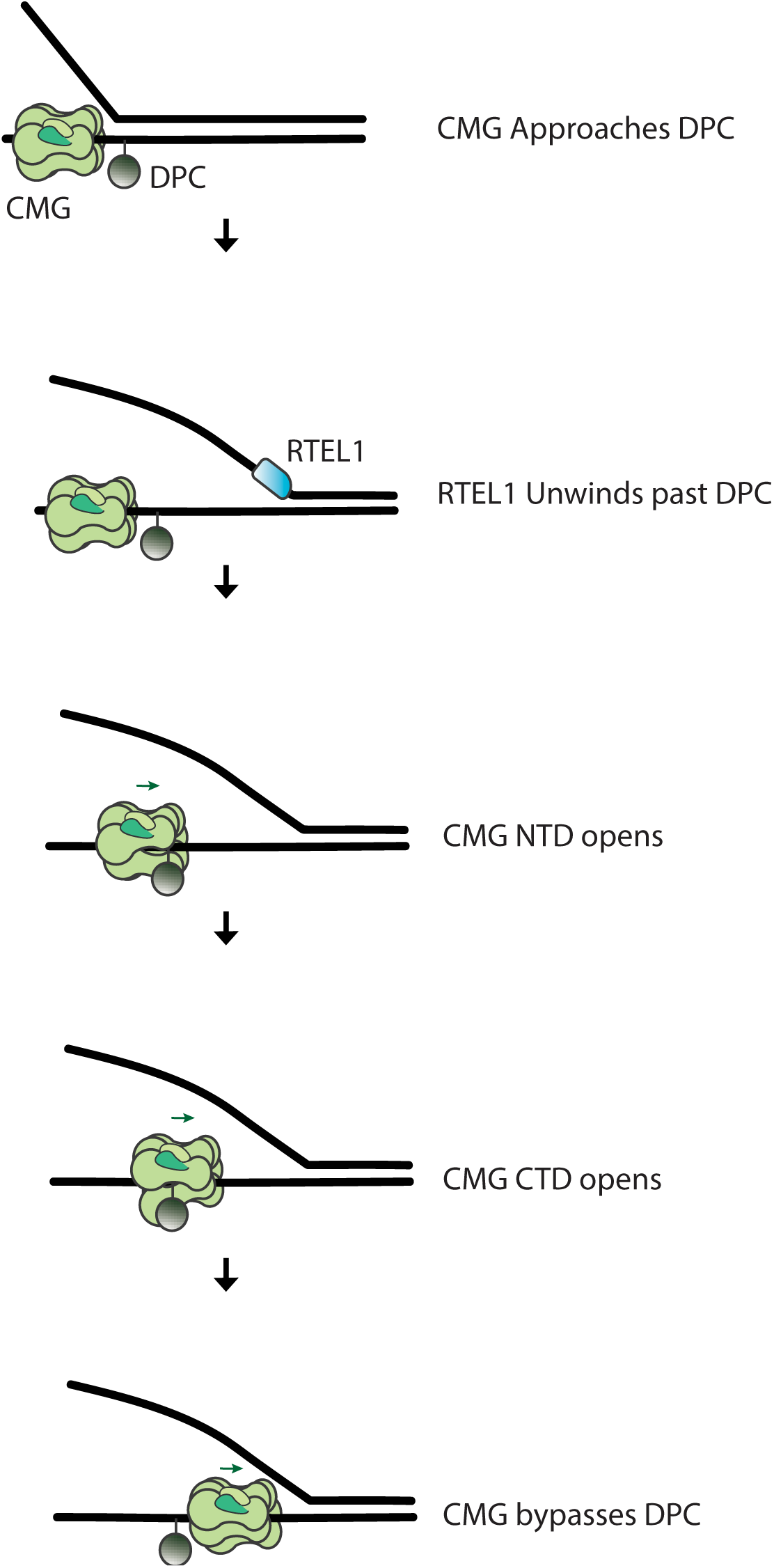
Model of DPC bypass by CMG. CMG initially stalls upon encountering a leading-strand DPC. RTEL1, acting on the lagging-strand template, unwinds past DPC and generates ssDNA downstream of the adduct. The leading N-terminal domain (NTD) of the MCM ring opens and grabs ssDNA available on the opposite side of the barrier. Once NTD closes the C-terminal domain (CTD) of MCM complex opens allowing the entire CMG complex to traverse the protein barrier.

### Versatility of the CMG helicase ring

The capability of the MCM ring to open is influenced by additional factors, which vary depending on the specific context of the replication process. During pre-RC assembly in yeast, for instance, Cdt1 is crucial for stabilizing the Mcm 2/5 interface opening [53], which is necessary for loading the Mcm complex onto the duplex DNA. Another key protein, Mcm10, is involved in the activation of CMG at a yeast replication origin, a step that includes transitioning the helicase from dsDNA to ssDNA [27, 59]. This requirement for Mcm10 may stem from its role in opening an Mcm interface in the yeast CMG complex, a hypothesis supported by the observation that the loading of recombinant pre-formed yeast CMG onto ssDNA is greatly enhanced in the presence of Mcm10 [27, 59]. Contrarily, at metazoan replication origins, MCM10 may not be as essential for helicase activation [60, 61]. Additional factors such as FANCM and DONSON have also been implicated in CMG ring opening, enabling the helicase to bypass DNA inter-strand crosslinks [62-64]. While the opening of *Drosophila* CMG during DPC traversal does not require any of the aforementioned factors in our reconstituted reactions, it remains conceivable that such proteins may enhance the ability of CMG to bypass DPCs in a cellular context.

Our findings indicate that the CMG complex has the capacity to open multiple MCM interfaces during the bypass of DPCs. This multi-interface flexibility may prove advantageous because, upon initial contact with a DPC, the obstructive protein could become wedged between the N-terminal Zinc finger domains of any two neighbouring MCM subunits. If the bypass mechanism was contingent on the opening of a unique MCM interface, CMG might necessitate a reorientation – potentially entailing a backward slippage along the DNA – until the DPC aligns against the appropriate gate. Although CMG has been observed to slide backward under certain conditions [65], such repositioning could be hindered by the interaction with the leading-strand polymerase epsilon. Moreover, this backward motion is typically enabled by the reannealing of the fork DNA, a process unlikely to occur if RTEL1 is actively unwinding the DNA downstream of the lesion. Consequently, we suggest that the CMG’s capability to use multiple MCM interfaces could be a strategic adaptation to effectively navigate past large DNA lesions.

### Preventing CMG disassembly during DPC bypass

The CMG helicase, being irreplaceable at an active replication fork, must be strictly safeguarded from dissociation. The interaction between CMG and the excluded strand at the fork junction is known to prevent helicase ubiquitylation and disassembly [50, 66, 67]. However, following RTEL1 unwinding the fork past a DPC, CMG should lose its engagement with the excluded strand, consistent with this interaction not being necessary for DPC bypass (Figure 5). This implies alternative mechanisms for protection against ubiquitylation during DPC traversal. Other replisome factors such as TIM/TIPIN are essential for tight association of the E3 ubiquitin ligase with CMG [66, 68]. One possibility is the dissociation of the TIM/TIPIN complex from CMG during bypass thus preventing association of the ubiquitin ligase. Notably, the release of TIM/TIPIN may also facilitate the opening of the MCM ring, as it binds across multiple subunits on the NTD of the MCM complex [69]. Furthermore, the structural reconfiguration of the MCM ring during DPC traversal may itself hinder ubiquitin ligase binding to CMG.

### Limitations of the CMG Helicase in Bypassing Large DPCs

While CMG is adept at bypassing a variety of DPCs, its ability to navigate past larger obstructions, such as SA, remains in question. It has been demonstrated that the FANCJ helicase unfolds DPCs to facilitate their proteolysis by the protease SPRTN [49]. Such unfolding by FANCJ is feasible post-CMG bypass, as FANCJ operates on ssDNA that is trailing the CMG, thus moving in the opposite direction. It has been speculated that FANCJ or a similar helicase could also initiate unfolding before CMG bypasses the DPC, potentially aiding the traversal of larger DPCs. However, there is a risk that such an unfolding helicase could collide with the CMG, potentially displacing it from the DNA. Hence, we consider that some DPCs, particularly larger ones, may be inherently insurmountable for CMG, requiring resolution through alternative mechanisms such as converging replication forks. Our findings in *Xenopus* egg extracts support this, as replication forks could not circumvent a covalent SA^Lead^, suggesting a potential limit to CMG’s capacity for bypassing substantial DPCs [23].

In summary, our work underscores the adaptability of the CMG complex in navigating replication stress, enhancing our understanding of its multifaceted role in promoting replication fork progression and genome stability.

## MATERIALS AND METHODS

### Generation of CMG variants

Various *Dm*CMG helicase constructs containing FRB, FKBP, DogTag, DogCatcher, or SnoopTag domains on individual MCMs were generated. Flexible linkers were added on either side of the inserted protein domains to ensure that both domains could interact on the surface of CMG. The linkers were copied from a previous study, which closed individual gates in the yeast Mcm complex [52]. The chosen insertion sites for the linkers were determined by aligning the yeast CMG (PDB: 5U8T) and *Drosophila* CMG structures (PDB: 6RAW). The protein domains including the linkers were cloned into pFastBac1 plasmids containing individual *Drosophila* MCM genes [17]. These were inserted after the following amino acid residue for each MCM (MCM2 – E194, MCM3 – D31, MCM4 – D190, MCM6 – E34, MCM7 – D25). MCM5 was tagged at the N terminus because internal insertions did not lead to CMG formation.

### Expression and purification of *Dm*CMG

All *Dm*CMG constructs were expressed and purified using previously described protocol with some modifications [23]. Each pFastBac1 plasmid carrying an individual *Dm*CMG subunit was transformed into DH10Bac *E.coli* competent cells, which serve as a host for the recombinant vectors, to generate bacmids. After bacmid generation, Sf21 insect cells were used for the initial transfection (P1 stage). We then conducted the next virus amplification stage to make P2 stocks using Sf9 cells. In all the virus amplification stages, cells were incubated in serum free Sf-900^TM^ III SFM insect cell medium (Thermo Fisher Scientific/Gibco) at 27°C while shaking at 120 rpm. To amplify the P3 stage of virus, 200 ml Sf9 cell cultures (0.5×10^6^ /ml) for each subunit were infected with 1 ml P2 viruses and incubated in 500 ml Erlenmeyer sterile flasks (Corning) for 4 days. Afterwards, P3 viruses for each subunit were mixed and used to infect 4 L of Hi5 cells (10^6^ /ml) with a multiplicity of infection of 5 (a total volume of 550 ml combined P3 viruses added to 1 L Hi5 cells). Approximately 6 L of infected Hi5 cells were divided into 500 ml aliquots using 2 L roller bottles (Corning) and incubated at 27°C for approximately 48 hours. Hi5 cells were harvested by centrifuging at 2000 ×g. Cell pellets were initially washed with PBS supplemented with 5 mM MgCl_2_, and then resuspended into 60 ml lysis buffer (25 mM HEPES pH 7.6, 1 mM EDTA, 1 mM EGTA, 0.02% Tween-20, 10% glycerol, 15 mM KCl, 2 mM MgCl_2_) with protease inhibitor cocktail tablets (Roche)) and flash frozen in liquid nitrogen in 10 ml aliquots before storing at -80°C.

All purification steps were performed at 4°C unless otherwise specified. The frozen cell pellets were first thawed in 28°C water bath and lysed via 60 strokes using a cell homogenizer (Wheaton, 40 ml Dounce Tissue Grinder) on ice. Next, 100 mM KCl was added drop wise and mixed into the cell lysate. Then the cell debris was removed via centrifugation at 24,000 ×g for 15 minutes. The supernatant was then collected and incubated with anti-flag M2 agarose beads (Sigma Aldrich) equilibrated with Buffer C (25 mM HEPES pH 7.5, 1 mM EDTA, 1 mM EGTA, 0.02% Tween-20, 10% glycerol, 1 mM DTT) for 2.5 hours. Next, agarose beads were washed 3 times with Buffer C-100 (25 mM HEPES pH 7.5, 1 mM EDTA, 1 mM EGTA, 0.02% Tween-20, 10% glycerol, 100 mM KCl, 1 mM DTT). CMG complex was eluted from the beads via incubation with Buffer C-100 (25 mM HEPES pH 7.5, 1 mM EDTA, 1 mM EGTA, 0.02% Tween-20, 10% glycerol, 100 mM KCl, 1 mM DTT) supplemented with 0.2-0.3 mg/ml DYKDDDDK peptide for 15 minutes at RT, followed by another round of incubation for 10 minutes at RT. The eluate two rounds were combined and filtered through a 0.22 µm Acrodisc syringe filter (PALL, #4650) and then passed through 1 ml HiTrap SPFF column (GE Healthcare) equilibrated with Buffer C-100. The sample was then separated via a 100-550 mM KCl gradient using Capto HiRes Q 5/50 column (GE Healthcare) across 60 ml. The fractions containing the CMG complex were pooled and dialyzed against CMG dialysis buffer (25 mM HEPES pH 7.5, 50 mM sodium acetate, 10 mM magnesium acetate, 10% glycerol, 1 mM DTT) overnight at 4°C. After dialysis, CMG was concentrated using a VivaSpin 500 concentrator with a 30,000 MWCO (Cytvia), aliquoted and flash frozen in liquid nitrogen before storing at -80°C.

### Fork DNA constructs

A variety of DNA substrates containing different modifications were designed. The sequences of oligonucleotides (oligos) used in each substrate can be found in Tables S1 and S2. Fork DNA substrates were created by combining equimolar amounts of required oligos (10 μM) in T4 DNA ligase buffer (50 mM Tris-HCl, 10 mM MgCl_2_, 1 mM ATP, 10 mM DTT, pH 7.5) from NEB. The oligo mixture was heated to 85°C for 2-5 minutes and slowly cooled down to room temperature (RT). For fork substrates requiring to seal ssDNA nicks, the oligo mixture was supplemented with 10 mM DTT, 2 mM ATP, T4 DNA ligase (NEB) and incubated overnight at RT. The reaction was later separated on 8% TBE PAGE in 1x TBE, the desired band was excised and placed in dialysis membrane (SpectraPor MW 3.5 kDa) containing TBE with 0.3 mg/ml BSA. DNA was purified from gel slice via electroelution in 1xTBE, gel slice was removed from the dialysis membrane, and DNA was dialyzed overnight at 4°C into buffer A (20 mM NaCl and 10 mM Tris pH 8.0).

To make DNA substrates containing site-specific leading- or lagging-strand methyltransferase DPCs, oligos were ordered to contain a 5-fluoro-2’-deoxycytidine (5FdC) modification. After purification, the modified fork at ∼1.5 μM was mixed with purified HpaII methyltransferase (M.HpaII) to 3.4 µM,, and supplemented with 100 μM S-adenosylmethione (SAM) in M.HpaII Buffer (50mM potassium acetate pH 7.9, 20 mM Tris-acetate, 10 mM magnesium acetate, 100 μg/ml BSA) and incubated overnight at 37°C. This reaction was then gel purified on an 8% TBE PAGE at RT to separate the DPC-modified DNA from the unmodified DNA. The bands corresponding to DPC-modified DNA were excised and isolated via electroelution, and subsequently dialyzed into buffer A. The relative concentration of fork DNAs were determined on a nanodrop (ND-1000 Spectrophotometer) and using a gel based serial dilution test scanned on a Fujifilm, SLA-5000 scanner. Finally, fork DNA was stored at 4°C.

Fork DNA substrates with thiol modification were created by combining equimolar amounts of oligos (10 µM) in phosphate buffered saline at pH 7.5 (PBS). The oligos were annealed and gel purified as before but dialysed overnight at 4°C into Buffer CJ (50 mM Na_2_HPO_4_/NaH_2_PO_4_, 150 mM NaCl pH 7.5). Alternatively, electroeluted fork DNA could be buffer exchanged into Buffer CJ with enough G50 columns equilibrated in Buffer CJ for the volume of fork DNA electroeluted. If the buffer exchange technique with G50 columns was used, the entire volume of fork DNA was exchanged through enough G50 columns at least twice. No amines can be present in the final reaction buffers, else downstream protein crosslinking does not work. If the volume of fork was greater than 150 µl after above steps, fork DNA was concentrated to at least 100 µl volume with a VivaSpin 500 concentrator with MWCOs of 3000 or 10000 (Cytvia). Buffer exchanged or dialyzed fork DNA substrate were then incubated with 0.5 µl of 0.5 M TCEP for 3 hours at RT or O/N at 4°C to reduce the thiol modification. One tube of commercial proFIre bifunctional crosslinking agent (Dynamic Biosensors) was then dissolved in 100 µl of water. Then, 50 µl of the dissolved bifunctional crosslinker was mixed with 100 µl of reduced fork DNA. The mixture was incubated for 5 minutes at RT. Next, G50 columns (Cytvia) were pre-equilibrated with Protein Conjugation Buffer (50 mM Na_2_HPO_4_/NaH_2_PO_4_, 150 mM NaCl pH 8.0) and 150 µl of the crosslinker-DNA mixture was buffer exchanged across the pre-equilibrated G50 columns (30 µl in each column). Buffer exchange was repeated on additional pre-equilibrated G50 columns. The combined buffer exchanged mixture was subsequently mixed with 50 µl of protein of interest at 5-10 mg/ml. The fork DNA and protein of interest mixture was then incubated for 1 hr at RT or O/N at 4°C. The reaction was later gel purified on an 8% TBE PAGE to separate DPC-modified DNA substates from unmodified DNA. The bands corresponding to DPC-modified DNA were excised and isolated through electroelution, and lastly dialyzed into Buffer A. The relative concentration of fork DNAs were determined on a nanodrop (ND-1000 Spectrophotometer) and using a gel based serial dilution test scanned using a Fujifilm, SLA-5000 scanner. These forks were also stored at 4°C.

### DNA unwinding assays

Approximately 75-100 nM of Cy5- or fluorescein-labeled DNA forks were mixed in Buffer B (20 mM NaCl, 10 mM Tris, 2 mM MgCl_2_, 0.3 mg/ml BSA) with 125 nM of CMG in CMG-binding buffer (25 mM HEPES pH 7.5, 5 mM NaCl, 10 mM Mg(OAc)_2_, 0.3 mg/ml BSA, and 1 mM DTT) and 125 µM ATPγS. This CMG/DNA mix was incubated at 37°C for 1 hr in a 7.5 µl total volume. Then 21 μl of ATP mix (CMG binding buffer with 1 mM ATP), containing excess polyT oligos (3 µM) and complementary oligos to the fork (0.3 µM), was added to 7 µl of the DNA-ATPγS-CMG mix to initiate unwinding. Excess oligos ensure reactions are single turn over by quenching free CMG, whereas complementary oligos ensure DNA does not reanneal behind CMG during unwinding and enhance CMG unwinding. Samples were then incubated at 30°C for varying amounts of time (30 sec to 1 hr) depending on the fork design and experiment. Reactions were terminated by the addition of 5 µl of stop buffer (0.2% SDS, 75 mM EDTA, 20% sucrose, 25 mM Tris pH 7.5). Reactions were then separated on an 8% TBE PAGE in 1xTBE. Gels were imaged on Fujifilm, SLA-5000 scanner using an appropriate laser and filter for each type of fluorescently labeled DNA fork. Cy5-labeled forks were scanned using 635-nm laser and Fujifilm LPR/R665 filter. Fluorescein-labeled forks were scanned using 473-nm laser and Fujifilm FITC filter.

Prior to starting the unwinding assay, using biotin-modified fork bound to monomeric streptavidin roadblock, the fork was diluted to ∼1 ng/µl into CMG buffer supplemented with 125 µM ATPγs. For each unwinding assay sample, 10 µl of diluted fork mixture was incubated with an additional 2 µl of monomeric streptavidin (Sigma SAE0094) dissolved to 0.05 mg/ml in conjugation buffer for 20 minutes at 37°C. The next step involved either loading CMG onto the fork DNA by the addition 1 µl of ∼150 nM CMG in CMG dialysis buffer or incubating with just CMG dialysis buffer for 1 hr at 37°C. To initiate unwinding, 6.5 µl of ATP mix (CMG binding buffer supplemented with 1 mM ATP, 3 µM polyT oligos, 0.3 µM complementary oligos to the fork, and 2.5 µM of the biotin containing oligo, oHYbio69) was added to the 13 µl of DNA/mSA/CMG mixture and then incubated at 30°C for various durations. Reactions were stopped by the addition of 5 µl of stop. The reactions were separated on an 8% TBE PAGE in 1xTBE and subsequently imaged on Fujifilm, SLA-5000 scanner using the appropriate laser and filter.

## ACKNOWLEDGEMENTS

We thank the past and present members of the Yardimci laboratory for critical and helpful discussions. This work was supported by the Francis Crick Institute, which receives its core funding from Cancer Research UK, the UK Medical Research Council and The Wellcome Trust (CC2133).

## Author contributions

M.T.K. and H.Y. designed the experiments. M.T.K. and E.d.L.P. cloned and purified various CMG constructs. S.X. expressed and purified most CMG constructs and performed unwinding assays using DogTag/DogCatcher CMG variants. M.T.K. performed all other experiments and analysed the data. M.T.K. and H.Y. wrote the paper with input from all authors.

