## Supplementary Figures for "Mechanism of DNA-protein crosslink bypass by CMG helicase"

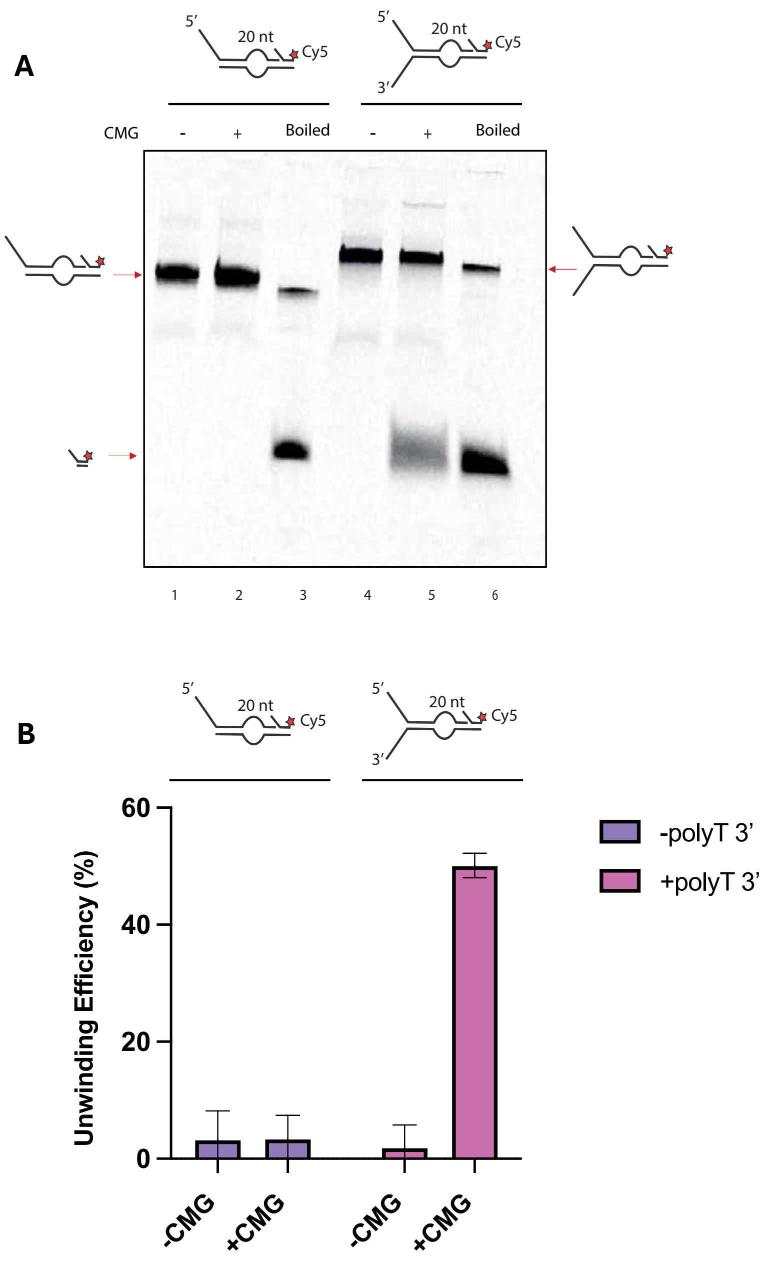


**Figure S1. CMG does not load onto the ssDNA bubble on model fork substrates.** Unwinding assay comparing fork DNA substrates containing or lacking a 3' polyT fork branch. All the forks contained a 20-nt ssDNA bubble followed by a downstream Cy5 labelled oligo on the excluded strand. **(A)** Example unwinding assay gel of the model fork substrates in the absence and presence of the 3' polyT fork branch. CMG was pre-bound to fork substrates in the presence of ATPγs at 37˚C for 1 hour. ATP and trap oligos were then added to initiate unwinding. Reactions were then incubated at 30˚C for 1 hour and stopped with SDS. The DNA was separated on an 8% PAGE and imaged for Cy5 fluorescence. **(B)** The quantification of unwinding efficiency of CMG on the various fork substrates, with the data represented as the mean ± SD from three independent experiments.

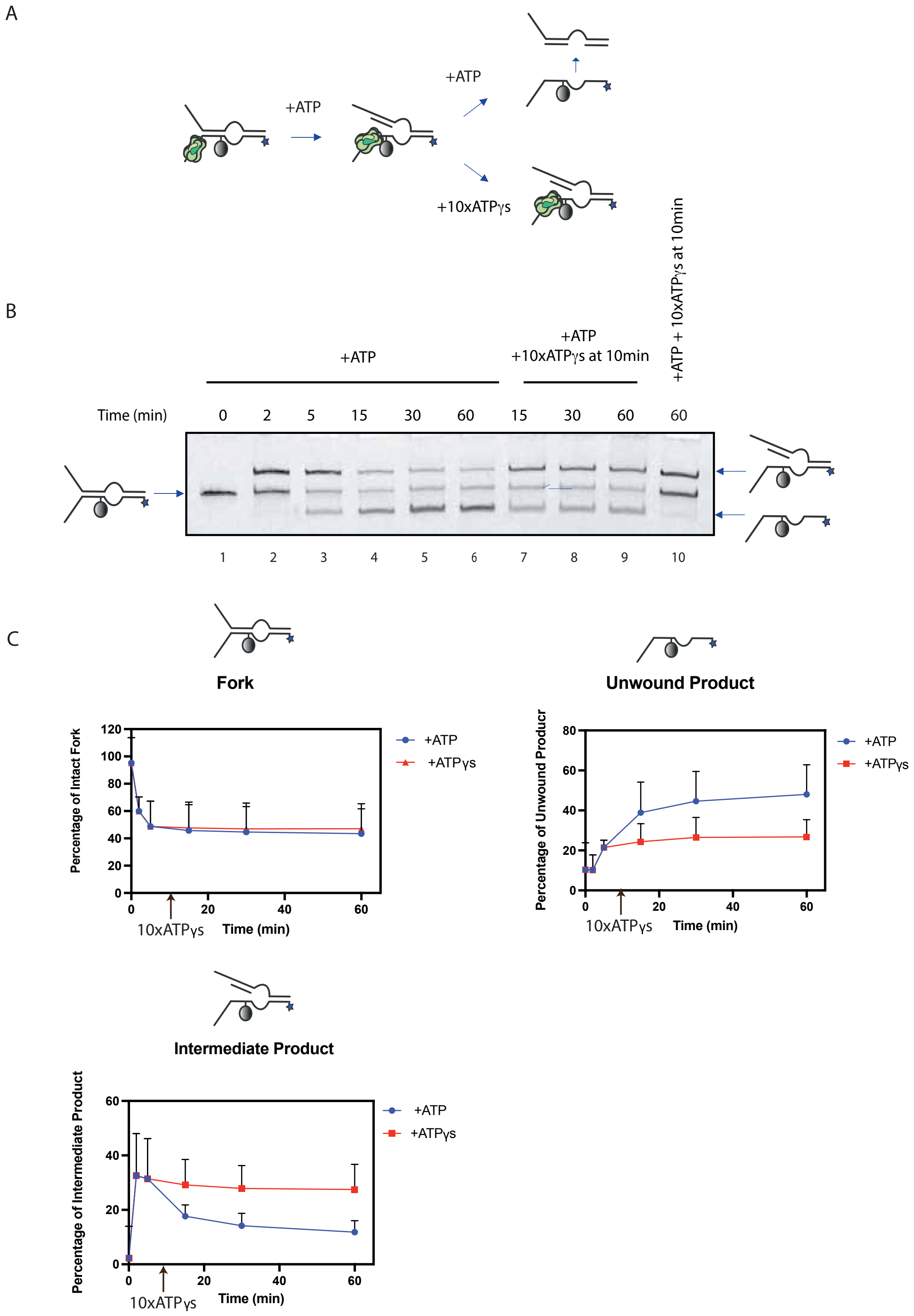
A
+ATP
+ATP
+10xATPγs
B
+ATP
+ATP
+10xATPγs at 10min
+ATP + 10xATPγs at 10min
Time (min)
0
2
5
15
30
60
15
30
60
60
1
2
3
4
5
6
7
8
9
10
C
Fork
Percentage of Intact Fork
120
100
80
60
40
20
0
0
20
40
60
10xATPγs
Time (min)
+ATP
+ATPγs
Unwound Product
Percentage of Unwound Producr
80
60
40
20
0
0
20
40
60
10xATPγs
Time (min)
+ATP
+ATPγs
Intermediate Product
Percentage of Intermediate Product
60
40
20
0
0
20
40
60
10xATPγs
Time (min)
+ATP
+ATPγs

**Figure S2. The unwinding of fork substrates into various unwinding products is CMG activity dependent.** After unwinding assay completion, a fork substrate can be separated into three components: the fork, an intermediate product that is partially unwound, as well as a fully unwound product. To show that CMG activity is required to form the various unwinding products a time course was conducted, but ATP was quenched during the time course with excess ATPγs. The fork substrate contained protein G as the leading-strand DPC as well as fluorescein on the translocating strand. **(A)** A cartoon diagram of the experimental design. The CMG is pre-bound to fork substrates in the presence of ATPγs at 37˚C for 1 hour. Then ATP and trap oligos were added to initiate unwinding. Reactions were incubated at 30˚C for various timepoints and stopped with SDS. However, at 10 minutes into the ATP incubation step, half the samples were further supplemented with a 10-fold excess of ATPγs over ATP, while the other half was not. Reactions were further incubated at 30˚C for various timepoints before stopping with SDS. DNA was then separated on an 8% PAGE and fluorescein fluorescence was imaged. **(B)** An example time course unwinding assay with the above experimental design and visualized on a native PAGE with fluorescence. **(C)** Quantification of the percentage of the various unwinding products (fork, unwound product, and intermediate product) overtime in the presence of absence of 10-fold excess ATPγs added at 10 minutes. The data are represented as the mean and SD from three independent experiments.

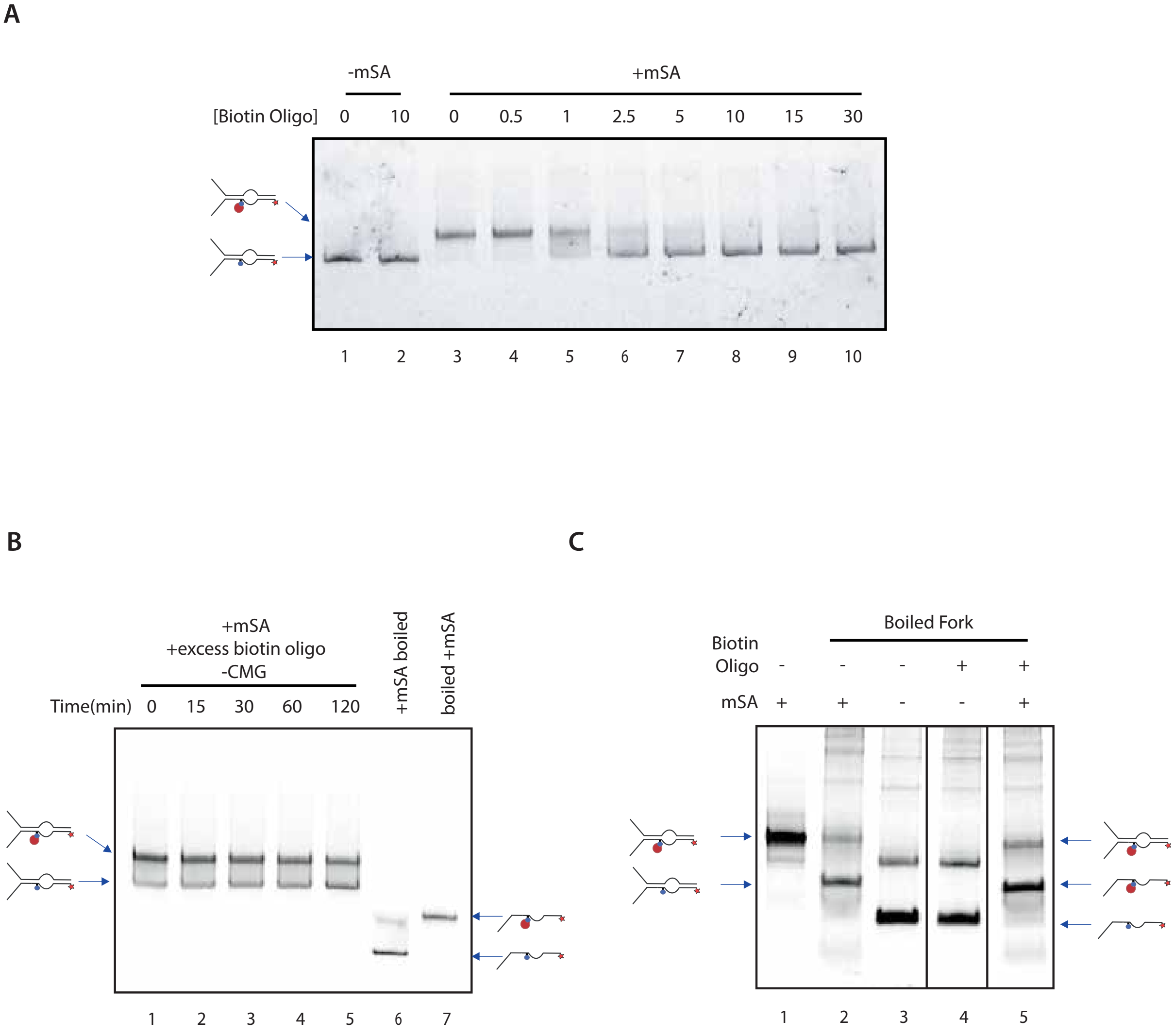


**Figure S3. Excess biotin prevents monomeric streptavidin binding to biotin-modified fork DNA.** Excess biotin was added by the incorporation of excess oligos containing a biotin-modification. **(A)** The biotin-containing Cy5-labeled fork used in the unwinding assays was incubated with monomeric streptavidin and increasing concentrations of biotin. At higher concentrations of biotin, free monomeric streptavidin cannot bind to the biotin-containing fork. **(B)** Biotin-modified fork was pre-bound to mSA, 2.5 mM of biotin was subsequently added and incubated for indicated length of time. The gel is visualized with Cy5 fluorescence showing that biotin does not significantly displace pre-bound mSA from biotin-containing fork substrate over time. **(C)** Heat denaturation test of the biotin fork with and without mSA. The test shows that competitor biotin does not displace mSA bound to heat-denatured fork substrate after a 2-hour incubation.

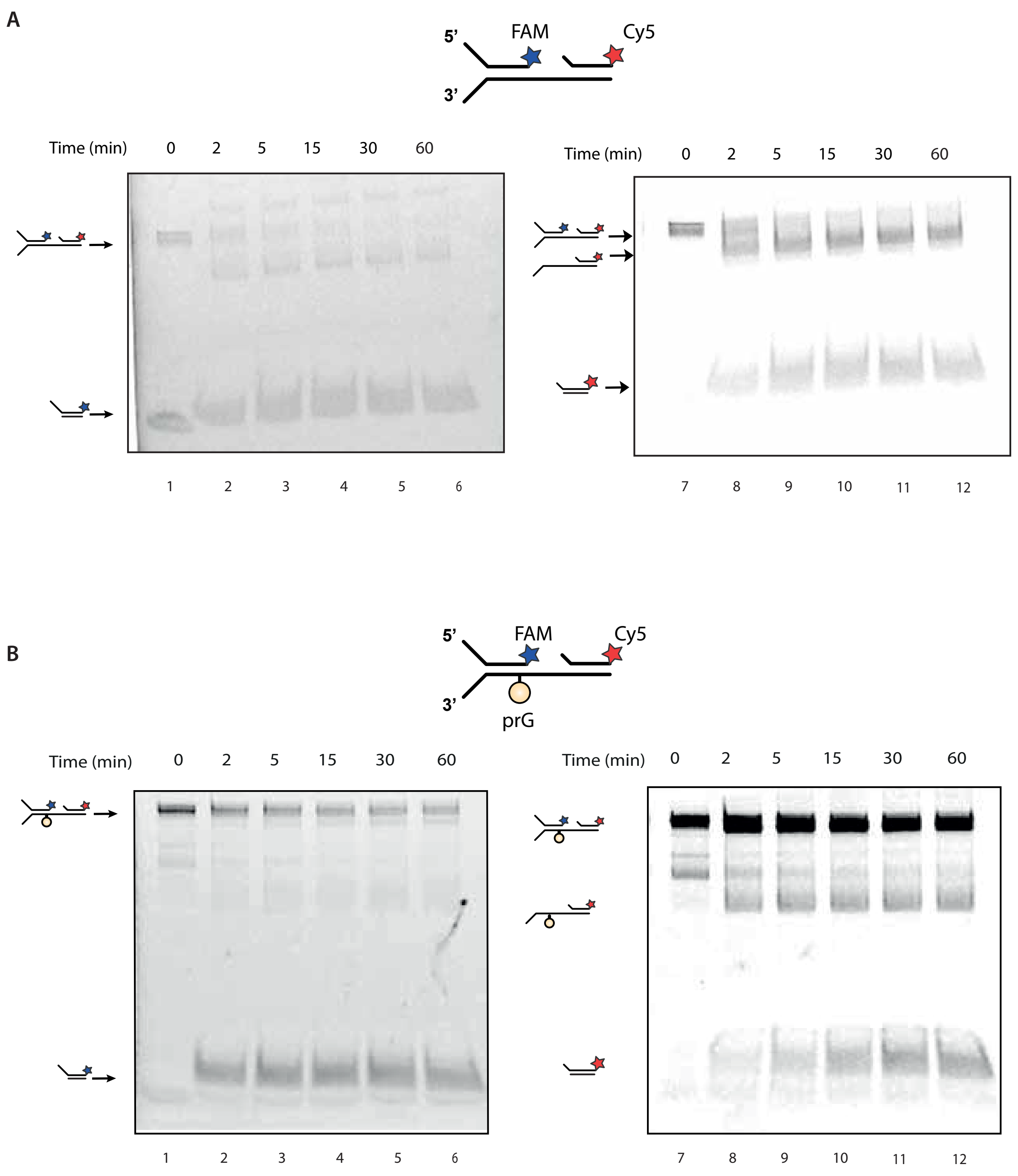


**Figure S4. CMG can bypass a DPC in the absence of the excluded strand.** Time course unwinding assay with the ssDNA gap carrying fork DNA substrates (A) without and (B) with a DPC. CMG was pre-bound to the fork substrates in the presence of ATPγs at 37˚C for 1 hour. ATP and trap oligos were then added to initiate unwinding. Reactions were further incubated at 30˚C for various durations and stopped with SDS. The DNA was separated on native PAGE, and subsequently both fluorescein (FAM) and Cy5 fluorescence were imaged. The left panels display fluorescein-labeled strand whereas the right panel displays Cy5 labeled strand.

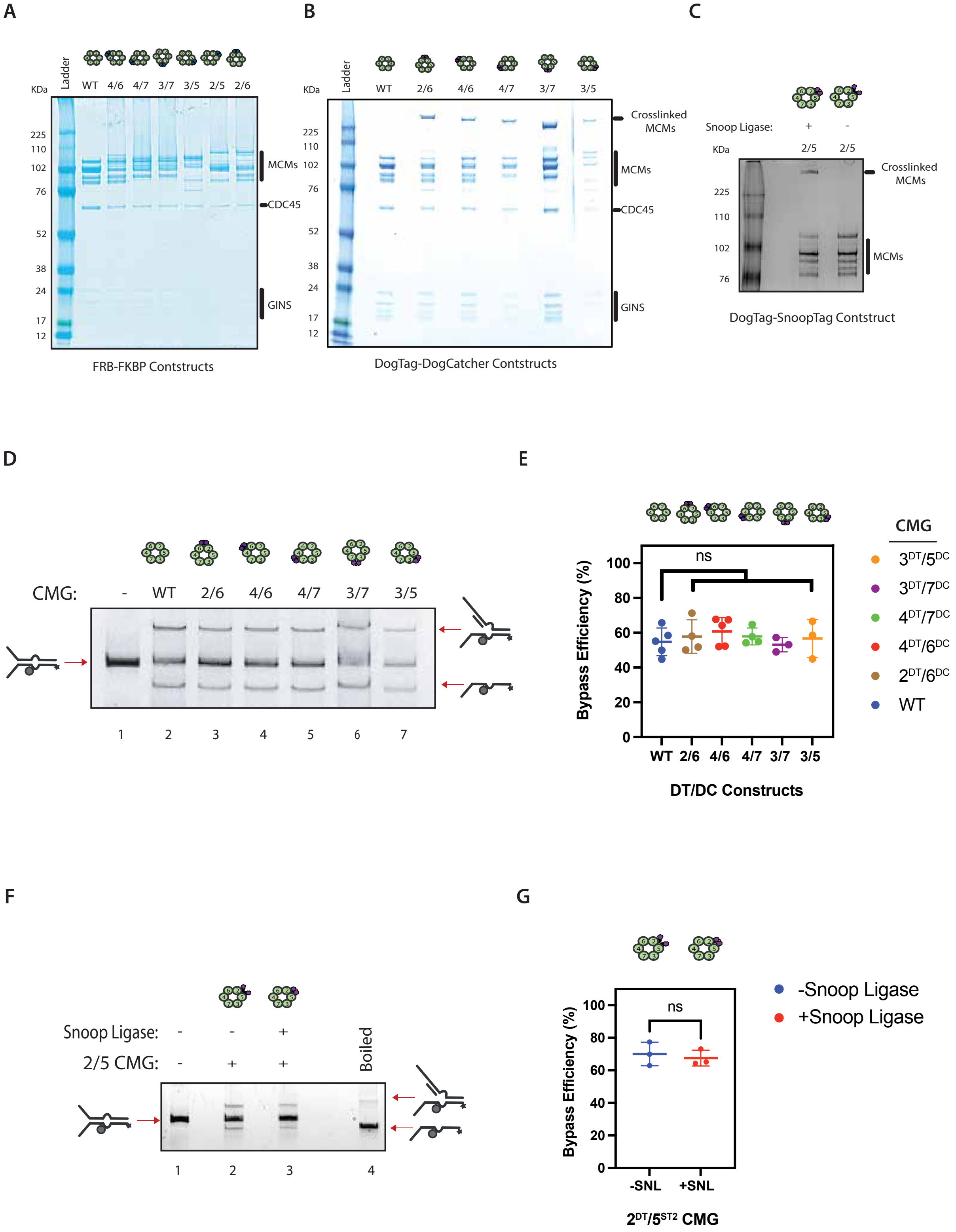
A
Ladder
KDa
WT 4/6 4/7 3/7 3/5 2/5 2/6
225
110
102
76
52
38
24
17
12
MCMs
CDC45
GINS
FRB-FKBP Contstructs
B
Ladder
KDa
WT 2/6 4/6 4/7 3/7 3/5
225
110
102
76
52
38
24
17
12
Crosslinked MCMs
MCMs
CDC45
GINS
DogTag-DogCatcher Contstructs
C
Snoop Ligase: + -
KDa
2/5 2/5
225
110
102
76
Crosslinked MCMs
MCMs
DogTag-SnoopTag Contstruct
D
CMG: - WT 2/6 4/6 4/7 3/7 3/5
1 2 3 4 5 6 7
E
Bypass Efficiency (%)
100
80
60
40
20
0
ns
WT 2/6 4/6 4/7 3/7 3/5
DT/DC Constructs
CMG
3DT/5DC
3DT/7DC
4DT/7DC
4DT/6DC
2DT/6DC
WT
F
Snoop Ligase: - - +
2/5 CMG: - + +
Boiled
1 2 3 4
G
Bypass Efficiency (%)
100
80
60
40
20
0
ns
-SNL +SNL
2DT/5ST2 CMG
-Snoop Ligase
+Snoop Ligase

**Figure S5. CMG variants to close individual MCM gates.** (A) A Coomassie-stained SDS polyacrylamide gel showing various purified CMG constructs each with two indicated MCM subunits modified to contain an FRB-FKBP pair. (B) A Coomassie-stained SDS polyacrylamide gel showing various purified CMG constructs each with two indicated MCM subunits modified to contain either DogCatcher (DC) or DogTag (DT) on an MCM pair. (C) A Coomassie-stained SDS polyacrylamide gel showing 2/5 CMG construct with MCM2 and MCM5 modified to contain DogTag (DT) and SnoopTag (ST2), respectively. Only, in the presence of the enzyme Snoop Ligase the two modified subunits will form an inducible covalent bond, effectively closing the MCM 2/5 interface, creating a larger MCM dimer. (D) Single-turnover fork DNA unwinding assays with various covalent closed gate CMG constructs. The fork substrate contained $MH^{Lead}$ followed by a 10-nt ssDNA bubble and a fluorescein (FAM) on the translocating strand. CMG was pre-bound to fork substates in the presence of ATPγs. ATP and trap oligonucleotides were then added to initiate unwinding. Reactions were further incubated at 30 ˚C for 1 hour and stopped with SDS. DNA was then separated on native PAGE and FAM fluorescence was imaged. (E) Quantification of DPC bypass efficiency of the different CMGs compared to wild-type (WT) CMG with data represented as the mean ± SD from three or more independent experiments. Unpaired T-test were performed to compare the bypass efficiency of each construct against the WT CMG; no modified construct was significantly different with $p > 0.05$ (ns) in all instances. (F) Single-turnover fork DNA unwinding assays with an CMG that contains DogTag (DT) and SnoopTag2 (ST2) on the MCM2 and MCM5 respectively. The MCM2 and MCM5 interface then can be covalently closed via the addition of the protein Snoop Ligase (SNL). The fork substrate contained $MH^{Lead}$ followed by a 10-nt ssDNA bubble and a fluorescein (FAM) on the translocating strand. CMG was pre-bound to fork substates in the presence of ATPγs. ATP and trap oligonucleotides were then added to initiate unwinding. Reactions were further incubated at 30 ˚C for 1 hour and stopped with SDS. DNA was then separated on native PAGE and FAM fluorescence was imaged. (G) Quantification of DPC bypass efficiency of the 2/5 mutant CMG in the presence of absence of Snoop Ligase with data represented as the mean ± SD from three independent experiments. Unpaired T-test was performed to compare the bypass efficiency of this construct with and without Snoop Ligase; $p > 0.05$ (ns).

**Table S1. Sequence of oligonucleotides used in this study.**

| Oligonucleotide Identifier | Sequence |
|---|---|
| oHY433 | GCTCTTTGTTCCTTCTCCTGTCCTTCCTTTGCTCGTTTTACAACGTCGTGCTGAGGTACCTTTTTTTTTTTTTTTTTTTTTTCTGTCCGGATGCTGAGTCAATGGGAATTCGTTTTTTTTTTTTTTTTTTTTTTTTTTTTTTTTTTTTTTTT |
| oHY442 | GACCTGCCTAGGAATTTTTTTATTCCTAGGCAGGTCCGAATTCCCATTGACTCAGCATCCGGACAGAATTTTTTTTTTTTTTTTTTTTGGTACCTCAGCACGACGTTGTAAAACGAGC |
| oHY443 | GCTCTTTGTTCCTTCTCCTGTCCTTCCTTTGCTCGTTTTACAACGTCGTGCTGAGGTACCTTTTTTTTTTTTTTTTTTTTTTCTGTCCGGATGCTGAGTCAATGGGAATTCGTTTTTTTTTTTTTTTTTTTTTTTTTTTTTTTTTTTTTTTT |
| oHY444 | GCTCTTTGTTCCTTCTCCTGTCCTTCCTTTGCTCGTTTTACAACGTCGTGCTGAGGTACCTTTTTTTTTTTTTTTTTTTTTTCTGTCCGGATGCTGAGTCAATGGGAATTCG |
| oHY449 | GACCTGCCTAGGAATTTTTTTATTCCTAGGCAGGTCCGAATTCCCATTGAC |
| oHYCy5_35 | GGCAGGCAGGCAGGCAGGCAGGCAGGCAGGCAGGCAGGCAGGAAGGACAGGAGAAGGAACAAAGAGC/3Cy5Sp/ |
| oHYCy5_38 | /5Cy5/GACCTGCCTAGGAATTTTTTTATTCCTAGGCAGGTCCGAATTCCCATTGACTCAGCATCCGGACAGAATTTTTTTTTTTTTTTTTTTTGGTACCTCAGCACGACGTTGTAAAACGAGC |
| oHYCy5_39 | /5Cy5/GCTCGTTTTACAACGTCGTGCTGAGGTACCTTTTTTTTTTTTTTTTTTTTTCTGTCCGGATGCTGAGTCAATGGGAATTCGTTTTTTTTTTTTTTTTTTTTTTTTTTTTTTTTTTTTTTT |
| oHYCy5_40 | /5Cy5/GACCTGCCTAGGAATTTTTTTATTCCTAGGCAGGTCCGAATTCCCATTGACTCAGCATCCGGACAGAATTTTTTTTTTGGTACCTCAGCACGACGTTGTAAAACGAGC |
| oHYCy5_43 | /5Cy5/GACCTGCCTAGGAATTTTTTTATTCCTAGGCAGGTCCGAATTCCCATTGACTCAGCATCCGGACAGAATTTTTGGTACCTCAGCACGACGTTGTAAAACGAGC |
| oHYCy5_52 | /5Cy5/GACCTGCCTAGGAATTTTTTTATTCCTAGGCAGGTCCGAATTCCCATTGACTCAGCATGGACAGAATTTTTTTTTTGGCTCTAGCAGGTACCTCAGCACGACGTTGTAAATGGAGC |
| oHYCy5Bio19 | /5BiotinTEG/GGCTCTAGCAGGTACCTCAGCACGACGTTGTAAATGGAGC/3Cy5Sp/ |
| oHYCy5Bio22 | /5Cy5/AGCTCCATTTACAACGTCGTGCTGAGGTACCTGCTAGAGCCTTTTTTTTTTT/iBiodT/TTCCATGCTGAGTCAATGGGAATTCGTTTTTTTTTTTTTTTTTTTTTTTTTTTTTTTTTTTTTTTT |
| oHYFAM14 | GACCTGCCTAGGAATTTTTTTATTCCTAGGCAGGTCTAGGGTCAGTTCGGTCCGATACACCATGACAT/36-FAM/ |
| oHYFAM15 | /56-FAM/AGCTCCATTTACAACGTCGTGCTGAGGTACCTGC |

| | |
|---|---|
| oHYFAM19 | /56FAM/GACCTGCCTAGGAATTTTTTTATTCCTAGGCAGGTCCGAATTCCCATTGACTCAGCATGGTTTTTTTTTTTTTTTGGCTCTAGCAGGTACCTCAGCACGACGTTGTAAATGGAGC |
| oHYFluo19 | GCTCGTTTTACAACGTCGTGCTGAGGTACCTTTTTTTTTTTTTTTTTTTTTTCTGTC/i5FdC/GGATGCTGA |
| oHYFluo20B | GCTCGTTTTACAACGTCGTGCTGAGGTACCTTTTTTTTTTTTCTGTC/i5FdC/GGATGCTGA |
| oHYFluo21 | /5Phos/TCAGCATC/i5FdC/GGACAGAATTTTTTTTTTTTTTTTTTTTGGTACCTCAGCACGACGTTGTAAAACGAGC |
| oHYFluo23 | GCTCGTTTTACAACGTCGTGCTGAGGTACCTTTTTTTCTGTC/i5FdC/GGATGCTGA |
| oHYFluo31 | GCTCCATTTACAACGTCGTGCTGAGGTACCTGCTAGAGCCTTTTTTTTTTTTTTTTTTTTTTTCTGTC/i5FdC/GGATGCTGAATGTCATGGTGTATCGGACCGAACTGACCCTATTTTTTTTTTTTTTTTTTTTTTTTTTTTTTTTTTTTTTTTTTTT |
| oHYTh3 | /5Phos/TAGAGCCTTTTTTTTTTTTTTCTGT/iDTPA/CCATGCTGAGTCAATGGGAATTCGTTTTTTTTTTTTTTTTTTTTTTTTTTTTTTTTTTTTTTTTTTTT |

**Table S2. Composition of DNA substrates and figures associated with each substrate.**

| Substrate Names | Oligonucleotides | Associated Figures |
|---|---|---|
| 20 nt ssDNA bubble Fork with MH Lead | oHYCy5_38, oHYfluo19, oHY433 | 1, 3 |
| 20 nt ssDNA bubble fork with MH Lag | oHYCy5_39, oHYFluo21, oHY449 | 1 |
| 20 nt ssDNA bubble control fork +polyT | oHY442, oHY443, oHYCy5_35 | S1 |
| 20 nt ssDNA bubble control fork -polyT | oHY442, oHY444, oHYCy5_35 | S1 |
| 10 nt ssDNA bubble fork with MH Lead | oHYCy5_40, oHYFluo20B, oHY433 | 6 |
| 5 nt ssDNA bubble fork with MH Lead | Cy5_43, oHYFluo23, oHY433 | 3 |
| 10 nt ssDNA bubble fork with Thiol DPCs | oHYCy5_52, oHYTh3, oHYFAM15 | 5, S2 |
| 20 nt ssDNA gap fork with MH Lead | oHYCy5Bio19, oHYFAM14, oHYFluo31 | 4, S4 |
| Biotin modified fork with 10 nt ssDNA bubble | oHYCy5Bio22, oHYFAM19 | 2, S3 |
